# Factors that influence the caste ratio in a bacterial division of labour

**DOI:** 10.1101/2024.02.28.582448

**Authors:** Luis Alfredo Avitia Domínguez, Zhengzhou Yu, Varun Chopra, Roeland Merks, Bram van Dijk, Daniel Rozen

## Abstract

Colonies of the bacteria *Streptomyces coelicolor* divide labour between cells that specialize on growth and sporulation and cells that specialize on antibiotic production. This division of labour arises due to costly chromosome deletions in the antibiotic overproducers. However, little is known about when and where these mutations occur or whether their frequency – which we liken to the caste ratio in social insects – is phenotypically plastic. To elucidate changes in the proportions of specialized cells (measured as the mutation frequency), we sampled *S. coelicolor* colonies grown under different conditions. Temporally, mutation frequency increased linearly with colony age and size. Spatially, mutations accumulated disproportionately in the colony center, despite greater growth and sporulation at the periphery. Exposing colonies to sub-inhibitory concentrations of some antibiotics, a competitive cue in *Streptomyces*, increased mutation frequencies. Finally, direct competition with other *Streptomyces* that naturally produce antibiotics increased mutation frequencies, while also increasing spore production. Our findings provide insights into the intrinsic and environmental factors driving division of labor in *Streptomyces colonies* by showing that mutation frequencies are dynamic and responsive to the competitive environment. These results show that chromosome deletions are phenotypically plastic and suggest that *Streptomyces* can flexibly adjust their caste ratio.

## INTRODUCTION

One of the hallmark features of multicellular organisms and complex societies is the division of labour among cells and individuals[1,2]. In multicellular species this leads to distinct cell types that perform different specialized functions[3]. Analogously, social animal groups divide into different castes that work together to increase colony efficiency[4]. In both cases, an assumption is that the proportion of cells or individuals performing specialized tasks is optimized to maximize group fitness[4–6]. This predicts that the caste ratio, measured as the level of investment into different castes, is not fixed but instead should vary according to the demands of the local environment.

Flexible caste ratios have been observed in several systems. Classic experiments with ants[7] and polyembryonic wasps[8] found that more soldiers were produced when colonies were exposed to higher levels of interspecific competition, and moreover that these shifts were adaptive. Similar results were recently reported in parasitic trematode worms that colonize snails as part of their complex life cycle [9,10]. As with ants and wasps, worms that developed in the presence of competition from other parasite species invested more heavily in defensive castes. This result supported the idea of flexible caste ratios while also broadening the range of species displaying this type of phenotypic plasticity. Our aim in this paper is to explore whether similar principles – that is to say, phenotypically plastic caste ratios– are also evident in multicellular bacteria.

*Streptomyces* are a group of multicellular bacteria with a complex life cycle[11,12]. Beginning as spores, colonies grow as an interconnected network of branching hyphae called the vegetative mycelium. When resources are exhausted, part of the colony differentiates to produce aerial hyphae that extend above the colony surface and ultimately give rise to a new generation of durable spores. This transition typically coincides with the production of secondary metabolites, including a diverse array of antibiotics[13,14]. *Streptomyces* are responsible for the majority of antibiotics used in medicine and agriculture in addition to an increasing variety of other compounds and enzymes[13,15].

The partition of the colony into vegetative and aerial hyphae results in a division of labour between growth and reproduction, respectively, that is equivalent to the germ-soma division in eukaryotes[11,16]. We recently discovered a second division of labour that acts solely within the vegetative hyphal population[17]. While most cells contribute to resource acquisition and growth, a minority of cells is responsible for producing metabolically expensive antibiotics[17,18]. An especially surprising aspect of this division of labour is the mechanism that drives it. In contrast to most bacteria, *Streptomyces* contain linear, rather than circular, genomes. It has long been known that these linear chromosomes are highly unstable, resulting in large deletions at the left and right chromosome flanks[19–22]. Deletions can be up to 1Mb and are invariably costly for the cells carrying them[18,19]. However, in the model species we study, *Streptomyces coelicolor*, strains with deletions hyper-produce a diversified set of antibiotics that these bacteria use to compete for food and space[18]. Through this mutation-driven division of labour[23], colonies circumvent the social conflicts associated with altruism, and can maximize both the production of spores and the production of antibiotics.

Although our previous work has clarified some of the costs and benefits of this bacterial division of labour, we know very little about the factors that give rise to this caste of antibiotic hyper-producing cells. Specifically, it is currently unknown i) when and where cells containing genome deletions arise during colony growth, ii) what factors regulate the fraction of cells containing such deletions, and iii) whether this fraction, *i*.*e*. the caste ratio, can respond flexibly to competition. Because competition in these bacteria is mediated by interference competition[24–27], specifically exposure to secreted antibiotics, we measured mutation frequencies following exposure to different antibiotics and then followed this with competition experiments in the presence of competing species that naturally produce antibiotics. While antibiotic exposure is a cue for competition[26,28,29], exposure to other species measures the direct response to both resource and antibiotic stress from a growing competitor. Briefly, we find that mutation frequencies shift as a function of colony size and age, while they disproportionately increase in the colony center. In addition, we observe significant changes in mutation frequencies in response to some antibiotics as well as a direct increase following exposure to competitors. These results reveal extensive plasticity in mutation frequencies and suggest that shifting the caste ratio can provide benefits when colonies face competitors.

## METHODS

### Bacterial strains and growth conditions

Three *Streptomyces* species were used in this study: *Streptomyces coelicolor* A3(2) (strain M145), *Streptomyces ardus* (DSM 110 40603) and *Streptomyces venezuelae* (DSM 40230). All strains were cultivated on soy flour mannitol medium agar plates (SFM) at 30°C. SFM contains, per liter: mannitol (20 g), agar (20 g), and soy flour (20 g). Spore stocks were prepared using standard protocols [30] and spore titers were determined by serial dilution.

### Mutation frequency through time

To study the temporal dynamics of mutations in *S. coelicolor* we diluted our frozen spore stocks to a concentration of roughly 50-100 spores/ml and then plated 100 ul on 80 separate SFM plates. This approach ensured that each plate contained 5-10 well isolated colonies whose growth was unimpeded by crowding. At nine different time points post-inoculation (72, 120, 168, 216, 264, 308, 360, 480 and 648 hours) we randomly sampled 16 well isolated colonies to determine CFU, mutant frequency and colony size, for a total of 144 colonies. Colonies with obviously aberrant morphology at the first three sampling points were excluded from sampling to avoid strains derived from mutated spores in the original spore stock.

To determine the size of each colony, pictures of the colonies were taken with a ZEISS AXIO Zoom V16 microscope for the first seven times and with a Nikon D200 Camera for the last two points due to their larger size. Colony area was measured in imageJ 1.54d version in which the dimensions of the colony were obtained based on pixel size.

Colonies were sampled by collecting the entire colony using a sterile loop and transferring the biomass to a 1.5ml microfuge tube containing 400μl of 30% glycerol. Spores were collected using a custom microfilter consisting of a cotton plugged 200μl pipette tip inserted into a hole drilled into the lid of a sterile 1.5 ml tube. Pre-assembled filters were covered in aluminum foil and autoclaved. Each colony was vortexed at maximum speed, after which 200 μl from the 30% glycerol solution was pipetted into the cotton-plugged pipet tip. Next, the microfilters were centrifuged at a speed of 12,000 rpm for 1.5 minutes which allowed spores to pass through while trapping the mycelia in the cotton. The process was finally repeated with the remaining volume (∼200 μl) from the original collection tube.

CFU was determined for each colony using serial dilution onto SFM agar and incubated for four to seven days at 30C. To ensure counts and mutation frequencies were obtained from well-dispersed colonies, we only used plates with 100-300 total colonies. Mutation frequency was estimated from the same plates by counting colonies with aberrant colony morphologies when compared to the wild-type strain. Mutant colonies had different coloration (indicative of altered antibiotic production or spore pigmentation), bald areas (producing no aerial hyphae), irregular shapes, notably smaller sizes, and/or no aerial hyphae development (Supplemental Figure 1). Putative mutants were independently scored by two people (LAD and ZY) and analyzed on the fourth day after sample inoculation. Mutation frequencies were quantified by dividing the number of mutants by the total CFU.

**Figure 1:**
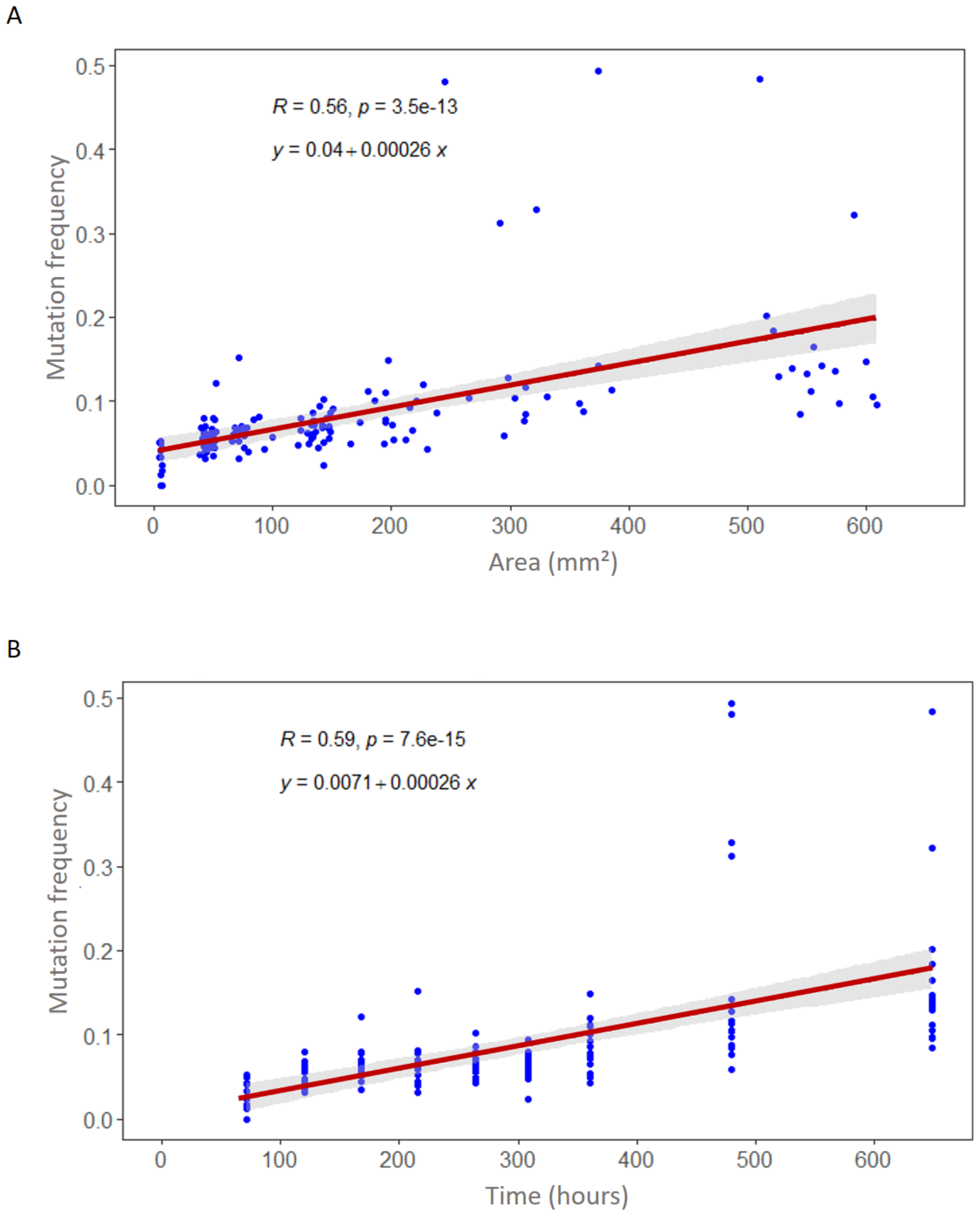
Mutation frequencies with respect of colony area (1A) and over time (1B).

### Spatial differences in mutant frequency

To assay the spatial distribution of mutants, colonies were plated as above onto SFM agar to ensure that there were roughly 5-10 well-isolated colonies per plate. Colonies (n = 32/time point) were sampled on Day 7 and Day 14 using the same custom microfilter as described above. However, rather than sampling the entire colony, we instead partitioned colonies into an inner and outer region (Figure 2). On Day 7, the interior of the colony was sampled by pushing the top of a sterile p200 pipette tip through the center of the colony and then depositing the biomass into a 1.5ml centrifuge tube containing 400μl of 30% glycerol, as above. The same procedure was used on Day 14, but with a larger p1000 tip. The outside of the colony for both time points was sampled with a loop and treated as above. The size of the sampled region at both time points corresponded to ½ the radius of the whole colony, leading to a roughly 4-fold difference in biomass area. In total, the area of the Day 7 colony corresponded to the interior area of the Day 14 colony, allowing a direct comparison between the two time points. CFU and mutant frequency were scored using the same procedure as above.

**Figure 2:**
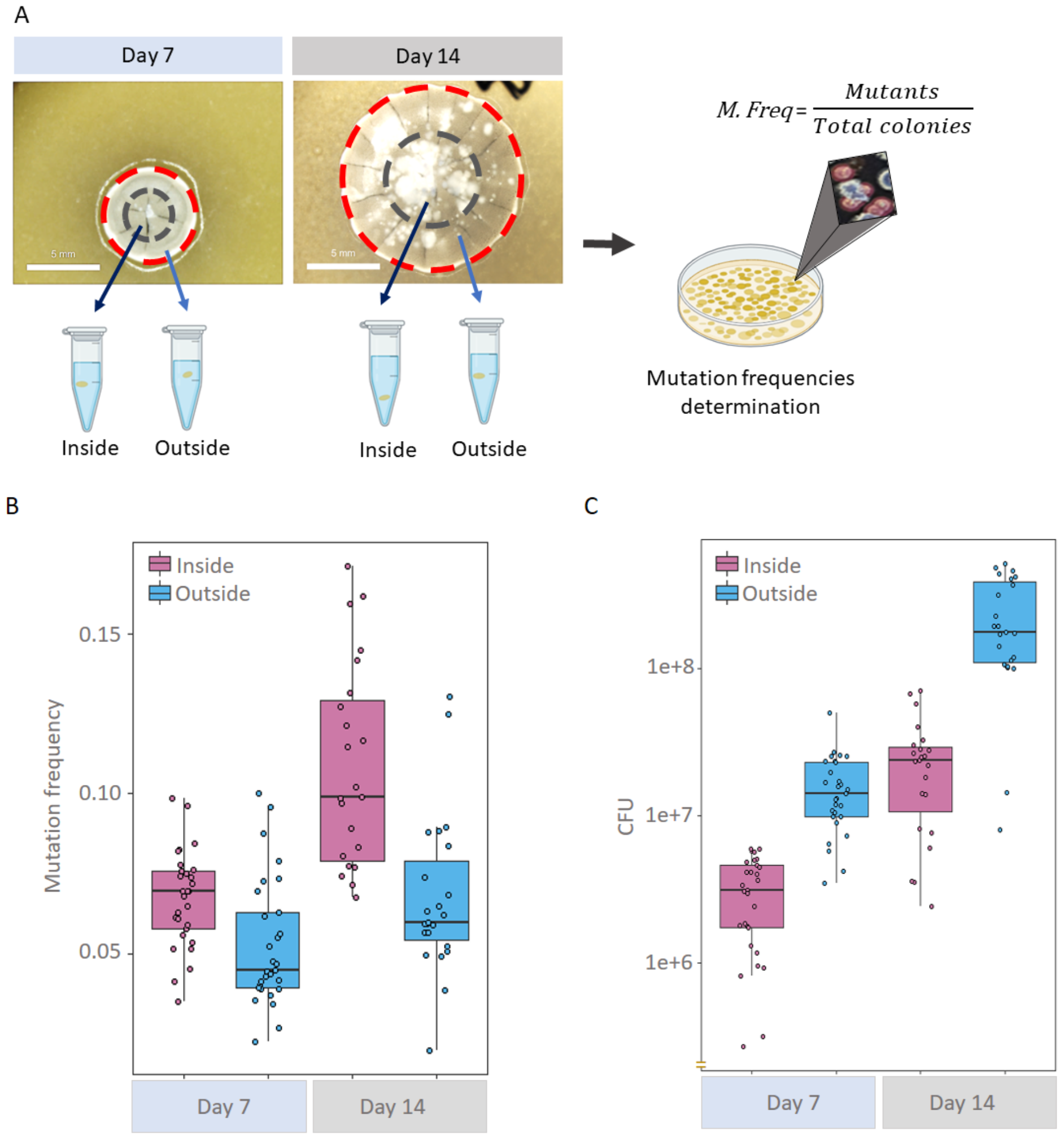
Spatial distribution of mutation frequencies in *Streptomyces* colonies. (A) Colonies were sampled from the interior (red circle) and exterior (gray line) on Day 7 and Day 14 days post inoculation. (B) Mutation frequencies of the interior (blue) and exterior (pink) portion of each colony at both sampling times: (C) CFU for the interior and exterior portions of the colonies.

### Effects of antibiotics on mutant frequency

To examine whether mutation frequency varied in response to antibiotic exposure, colonies of *S. coelicolor* were grown in the presence of sub-MIC concentrations of chloramphenicol, erythromycin, gentamycin, kanamycin, streptomycin, tetracycline, rifampicin, mitomycin and novobiocin. Antibiotics were chosen based on prior experiments from our own preliminary studies and work from other labs, and also to include antibiotics with different modes of action. MIC for each antibiotic was determined by plating a spot of 10^5^ spores on plates with different antibiotic concentrations; MIC was scored as the lowest antibiotic concentration that inhibited growth. Mutation frequencies were assayed at 25% of the MIC for all drugs except for streptomycin, where we used 10% of the MIC to avoid effects on viability. The MIC for each antibiotic is shown in Table 1. Mutation frequencies and CFU for each colony were determined as described above.

**Table 1.** MICs of different antibiotics to *S. coelicolor* growing on SFM agar. Note: CMP – chloramphenicol, ERY – erythromycin, GEN – gentamycin, KAN – kanamycin, STR – streptomycin, TET – tetracycline, RMP – rifampicin, MMC – mitomycin C, NB – novobiocin.

| Antibiotic | CMP | ERY | GEN | KAN | STR | TET | RMP | MMC | NB |
| --- | --- | --- | --- | --- | --- | --- | --- | --- | --- |
| MIC ($\mu\text{g/ml}$) | 42 | 8 | 10 | 5 | 2 | 30 | 10 | 2 | 3 |

### Effects of interspecific competition on mutant frequency and CFU

To test if mutation frequencies were increased due to interactions with competitors, we established co-culture experiments between *S. coelicolor* and two other species, *S. venezuelae* and *S. ardus*. Both species naturally produce antibiotics that were found to increase mutation frequencies (Figure 3). *S. venezuelae* produces chloramphenicol while *S. ardus* produces mitomycin C. In the co-culture, the focal strain, *S. coelicolor*, and competitor strains, *S. ardus* and *S. venezuelae*, were streaked separately on SFM agar along the opposite edges of 1cm X 1cm wells of a 25-well plate. The concentration of the spores was adjusted so that there were approximately 20-30 competitor colonies and two to four *S. coelicolor* colonies. The distance between competitors was approximately 8mm. Colonies were plated simultaneously or with a two-day delay to test if increased antibiotic secretion from the competitor would further increase the mutation frequency. Well separated colonies were sampled after 7 days of growth using the protocol above.

**Figure 3:**
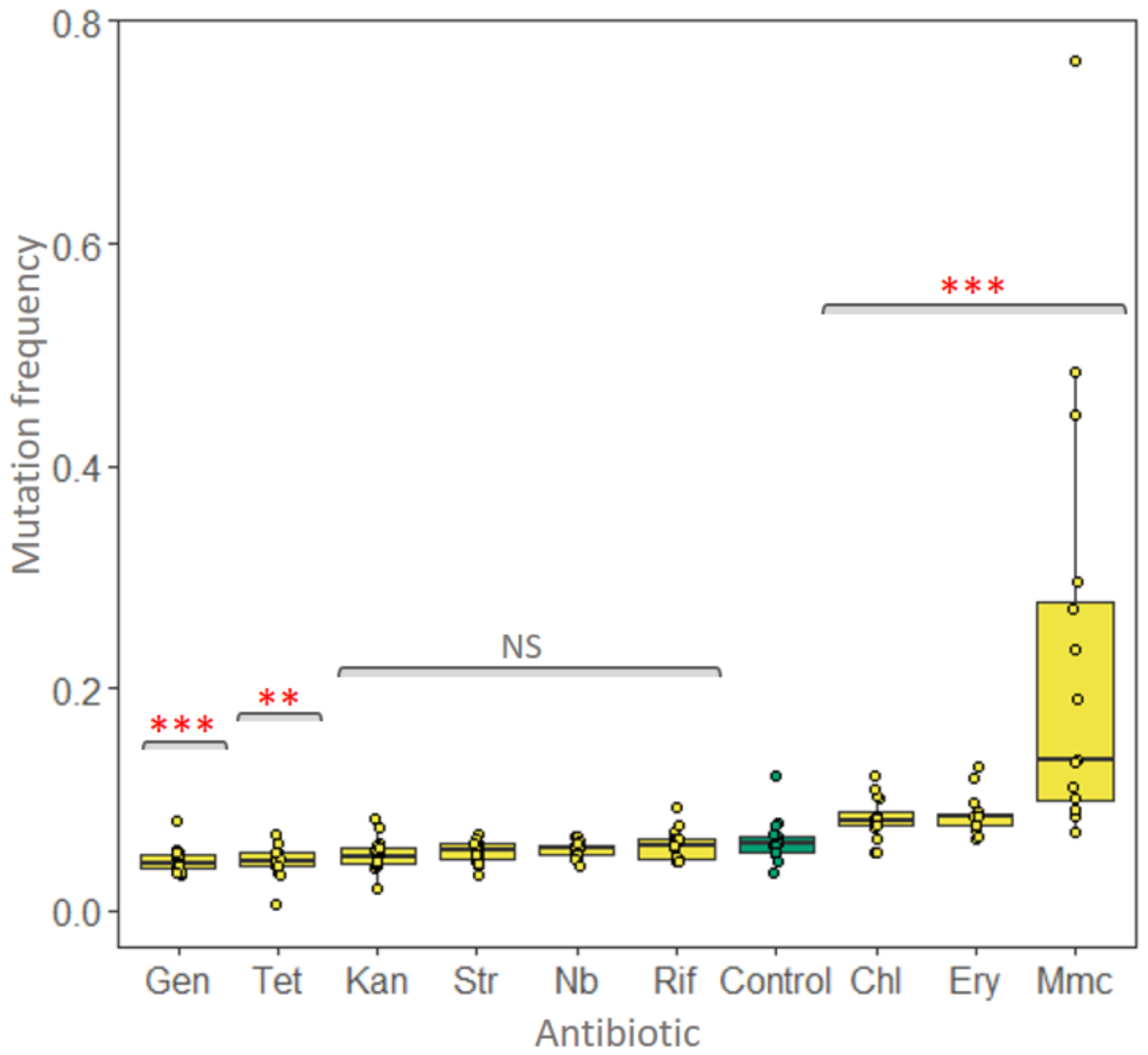
Mutation frequencies of *S. coelicolor* grown in the presence of ¼ MIC of different antibiotics. Tetracycline = Tet, Gentamicin =Gen, Kanamycin = Kan, Streptomycin = Str, Noboviocin = Nb, Rifampycin = Rif, Erythromycin = Ery. Mitamycin = Mmc were used. Control colonies grown without antibiotics are shown in green. NS = not significant, p < 0.01 = **, p < 0.001 = ***.

### Statistical analysis

Statistical analysis was performed using the *stats* package of R studio Software V 4.0.4.[31] Linear regression was performed using the *lm* function to determine the relationship between of mutation frequency, time and colony area or CFU. Tests of significance were conducted to evaluate the effect of the different treatments in each assay. Because data were not normally distributed (via Shapiro-Wilkinson tests), mutation frequencies were analyzed using Mann-Whitney U tests. Wilcoxon signed-rank tests (V) were used to compare paired mutation frequencies in the interior and exterior of bacterial colonies. Additionally, a one-sample Wilcoxon signed rank test (W), with a mu value of one, was used to test the ratio of interior vs exterior mutation frequencies. *ggplot* V 3.3.5, *ggpubr* V 0.4.0, and *tidyverse* V 1.3.1 packages were used to generate graphs.

## RESULTS

### Temporal dynamics of mutation

To track changes in mutation frequency through time, we destructively sampled a total of 144 colonies taken every few days after growth on SFM agar plates. Our results in Figure 1 show that mutation frequency increases significantly as colonies grow and age (area: R = 0.56, p < 0.001; age: R = 0.59, p < 0.001). In extreme cases, mutation frequencies can approach 50%. As expected, both colony area and CFU increased continuously through time and are significantly correlated (R = 0.78, p < 0.001) (Supplemental Figure 2).

### Spatial dynamics of mutation

Data and theory suggest that mutations that arise at the colony edge, where most growth occurs, are more likely to fix via the process of “allele surfing”. To test if results in Figure 1 could be explained by this possibility we quantified mutation frequencies and CFU from the center and the edge of 7 and 14 day-old colonies. As expected, CFU and mutation frequency both increased from Day 7 to Day 14 (CFU: U = 34, p < 0.001 ; mutation frequency: U = 52, p < 0.001). While the CFU and mutation frequency on Day 7 were ∼9 x 10^6^ and ∼0.06 on average, they were 1 x 10^8^ and ∼0.09 on Day 14. Looking more closely, however, we observed striking differences as a function of where spores were sampled. CFU between Day 7 and Day 14 increased almost exclusively at the colony edge. That is to say, the spore numbers in the interior of Day 14 colonies did not differ from the number of spores in Day 7 colonies (U = 277, p = 0.305), while we observed a significant CFU increase on the colony edge (U = 45, p < 0.001). The situation is reversed for mutation frequency. While the mutation frequency on the inside of the colony was significantly increased in Day 14 colonies (U = 45, p < 0.001), the outside remained unchanged (U = 256, p = 0.15). On both Day 7 and 14, the mutation frequency was significantly higher in the inside vs the outside of the colony (paired Wilcoxon signed rank test: Day 7: V = 361, p = 0.001; Day 14: V = 263, p < 0.001). However, there were no significant differences in the ratio of interior:exterior across time points (W = 262, p = 0.19). Thus, although our data showed evidence of task division between the interior and the periphery of *Streptomyces* colonies, the spatial location of the castes was the reverse of predictions.

### Antibiotic responsiveness

To determine if mutation frequency was influenced by external conditions, i.e., to test if external conditions could give rise to shifts in caste ratios, we exposed colonies of *S. coelicolor* to sub-minimum inhibitory concentrations (sub-MICs) of a set of diverse antibiotics. Antibiotics are ecologically relevant cues in *Streptomyces* as these species are prolific producers of these anti-competitor toxins. Of nine tested compounds, five caused mutation frequencies that were significantly different from what we observed on drug-free agar (Figure 3). Of these, three were higher than the control (chloramphenicol, erythromycin and mitomycin C) and two were lower than the control (gentamicin and tetracycline). We observed no difference in mutation frequencies for cells exposed to kanamycin, streptomycin, rifampicin or novobiocin (see figure legend for statistical details). These results show that environmental cues (antibiotics) that are naturally produced by competing *Streptomyces* species can significantly modify the caste ratio in *S. coelicolor* colonies.

### Flexible caste ratios in response to competition

As shown above, several antibiotics can induce increased mutation when colonies are grown on plates containing sub-MICs of antibiotics. However, the antibiotic concentrations we used during these assays are not necessarily reflective of the concentrations *Streptomyces* colonies experience during direct interspecific competition. We therefore asked if direct competition with *S. venezuelae* and *S. ardus* which naturally produce chloramphenicol and mitomycin C, respectively, sufficed to induce mutations in *S. coelicolor*. Both antibiotics caused an increased mutation frequency in *S. coelicolor* (Figure 3).

As shown in Figure 4a, in pairwise competition assays we found no difference between the mutation frequency of *S. coelicolor* when it was grown alone and when it was grown in competition with another *S. coelicolor* colony (U = 79, p = 0.98). As expected, CFU declined as a consequence of resource competition between adjacent colonies (Figure 4B) (U = 160, p < 0.001). When *S. coelicolor* was grown adjacent to *S. ardus*, we also observed no change in mutation frequency when cells were plated simultaneously (U = 35, p = 0.70). However, when we allowed *S. ardus* to grow for two days prior to plating *S. coelicolor*, mutation frequency was significantly increased (U = 2, p < 0.001) (Supplemental Figure 3), suggesting that secretion of mitomycin C increased over time. Similarly, mutation frequency was significantly increased when *S. coelicolor* was grown adjacent to *S. venezuelae* (U = 5, p < 0.001). However, in contrast to the response to *S. ardus*, mutation frequency already increased during simultaneous exposure to *S. venezuelae*, while further exposure (for two days) had no additional effect (U = 54, p = 0.73). Importantly, the CFU of *S. coelicolor* increased during competition with *S*.*ardus* and *S. venezuelae* (U = 14, p = 0.02; U = 11, p = 0.009) as compared to the control when it competed against itself. Thus suggests that the increased mutation frequency may provide a competitive advantage through increased antibiotic production.

**Figure 4:**
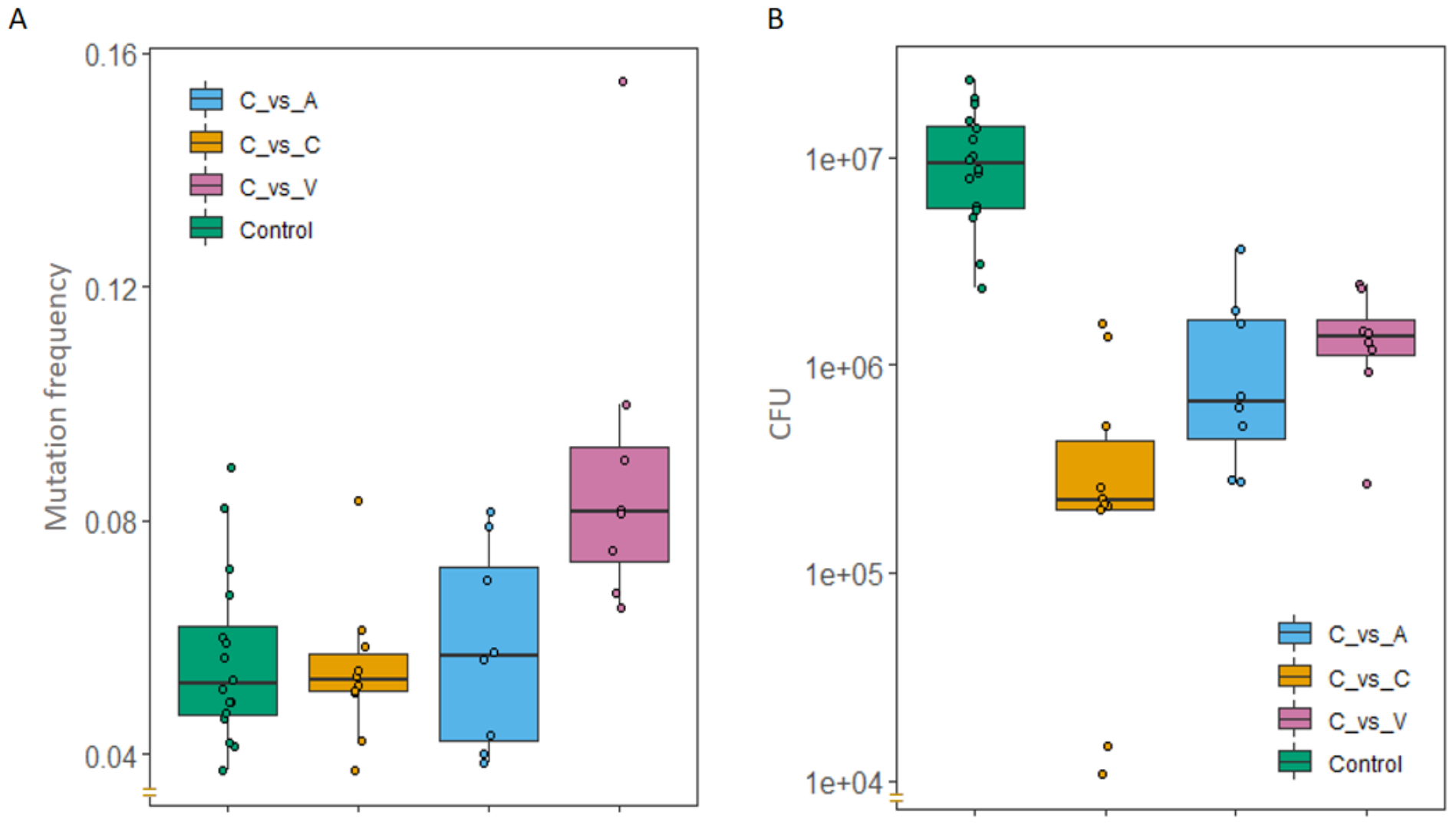
Competition between *Streptomyces coelicolor* and two other species of *Streptomyces*. C = *S. coelicolor*, A = *Streptomyces ardus*, V = *S. venezuelae*. Control treatments correspond to *S. coelicolor* grown in the absence of a competitor.

## Discussion

The linear chromosomes of Streptomyces have long been known to be unstable at the chromosome ends[19,21,22]. While the molecular mechanisms behind this phenomenon remain unclear, large chromosomal deletions are of considerable interest to industry because they potentially threaten the stability of antibiotic or enzyme production in industrial fermenters[32]. We recently studied the ecological and evolutionary consequences of chromosome instability in *S. coelicolor* and found that these mutations gave rise to a division of labour, whereby mutated cells overproduce antibiotics for the rest of the colony[17]. However, it remained unknown if the fraction of cells with mutations, which is analogous to the caste ratio of social insects, varied in different conditions or stages of growth.

Here, we quantified mutation frequencies of hundreds of independent colonies through time and in different regions of the colony. Our results provide clear evidence that the mutation frequency increases as colonies grow and age (Figure 1). Previous studies with unicellular bacteria have obtained similar results and hypothesized that this is due to the accumulation of e.g. reactive oxygen species (or other compounds) that are mutagenic or represses DNA repair[33,34]. However, an increased frequency of mutants does not necessarily imply that the basal mutation rate has also increased. “Allele surfing”, where mutants have an increased probability of fixation at the colony edge due to increased resource access, can also lead to a similar outcome without any change in mutation rate[35,36].

To distinguish these possibilities, we dissected colonies by separately analyzing CFU and mutation frequencies from the middle and periphery of the colony (Figure 2). As expected in growing colonies, CFU increased significantly on the edge of the colony between Day 7 and Day 14. However, in contrast to the predictions of “allele surfing”, mutation frequency only significantly increased in the middle of the colony, despite the fact that CFU remained unchanged in this region. One possible cause of this result is if chromosome deletions occurred in non-replicating spores due to exposure to mutagenic compounds (e.g. antibiotics) that are predominantly produced in the middle of the colony. Alternatively, there may be continued growth, chromosome replication and sporulation in the center of the colony, but this is offset by germination of older spores, leading to no net change in CFU. Older studies relying on detailed microscopy confirm extensive “precocious” germination within colonies that could explain this dynamic[37]. However, increased mutation frequencies would then require that newly formed spores have a higher mutation rate. *S. coelicolor* produces several developmentally regulated antibiotics, including the blue-pigmented actinorhodin and the red-pigmented undecylprodigiosin[14,15]. While actinorhodin has no known mutagenic functions, prodigiosin is a known mutagen whose expression is increased in aging hyphae[38–41]. These features make it a strong candidate for increasing mutation frequency in aging colonies and in a spatially restricted manner. We intend to test this directly in future studies using strains lacking prodigiosin and in environments where we can disrupt the spatial localization of this compound.

While intrinsic factors like prodigiosin might unavoidably increase mutation rates in aging colonies, external factors could potentially adaptively increase mutation frequencies. Antibiotics are an ecologically relevant environmental cue for *Streptomyces* given the role of these metabolites in interference competition[15,24,27,42]. Moreover, exposure to antibiotics, either with fixed concentrations in agar[43] or during co-culture with a competing strain[26,28], can induce antibiotic production. As shown in Figure 3, several antibiotics significantly shifted the mutation frequency; however, we observed both increases and decreases, even with antibiotics with similar modes of action on protein synthesis. This suggests that antibiotics may not be a general driver of increased mutation in *Streptomyces*, but rather that this depends on the specific mode of action or other unknown factors. Additionally, these assays may be sensitive to the antibiotic concentration we used on our assays. Preliminary data with three antibiotics (novobiocin, ciprofloxacin and mitomycin C) (VC, unpublished data), as well as results from other groups[44–46], show a strong dose dependence on mutation rates. Higher concentrations markedly increase mutations, but they also dramatically reduce CFU. It was for this latter reason that we chose to analyze strains at ¼ MIC.

Because of potential limitations of using arbitrary concentrations of antibiotics, we carried out direct competition assays between species that naturally produce chloramphenicol and mitomycin C (Figure 4). In both cases, we observed an increased mutation frequency, although this only occurred after giving *S. ardus* a two-day head start to allow sufficient accumulation of mitomycin C. More interestingly, this change coincided with increased spore production, suggesting that a higher fraction of mutant strains within colonies led to increased fitness. As yet, it remains unknown if this adaptive response is due to the change in mutation frequency, to antibiotic induction due to competition sensing, or some combination of the two. Regardless, this supports the idea that increased antibiotic production in *S. coelicolor*, whether via mutation[17] or gene regulation[47], is phenotypically plastic and can respond adaptively to competition with other species.

The specific aim of this paper was to analyze intrinsic and extrinsic factors regulating mutation frequencies, and therefore the expression of division of labour, in *S. coelicolor*. We use mutation frequency as a proxy for social caste in analogy with socially determined castes in colonies of insects or other animals. While we believe our results uncover ecologically relevant plasticity in mutation rates in *S. coelicolor*, there are limitations to our conclusions and to the analogy with social insects. First, our determination of “caste” is limited to strains with obviously aberrant morphology, which may lead to a systematic underestimate of caste ratios if some fraction of chromosome deletions do not cause phenotypic changes. Second, as yet, the causal link between increased mutation frequency and increased competitiveness remains unclear. We know that a higher mutant fraction within colonies leads to higher antibiotic production, but further work will be required to determine how this results in increased CFU during competitive interactions. Finally, even though the present work shows that the caste ratio in *Streptomyces* is “flexible”, it remains unknown if it can be optimized to ensure the greatest benefit from antibiotic production (due to increased mutation rate) at the lowest cost to the colony. Optimal caste ratios have been extensively studied in social insects[6], but there is no comparable theoretical literature on bacterial divisions of labour. Addressing these experimental and theoretical limitations remains an important objective of future work.

## Acknowledgements

We thank Michael and Barbara Taborsky for inviting us to join this special issue and for organising the “Division of labour as key driver of social evolution” workshop at the Wissenschaftskolleg zu Berlin in March 2023. DER would like to acknowledge funding from the NWO (OCENW.M.22.076) and the DFG-SPP2389 (TS 325/4-1).

## Notes

### Competing Interest Statement

The authors have declared no competing interest.

