## Supplemental files for "Factors that influence the caste ratio in a bacterial division of labour"

Supplementary Material

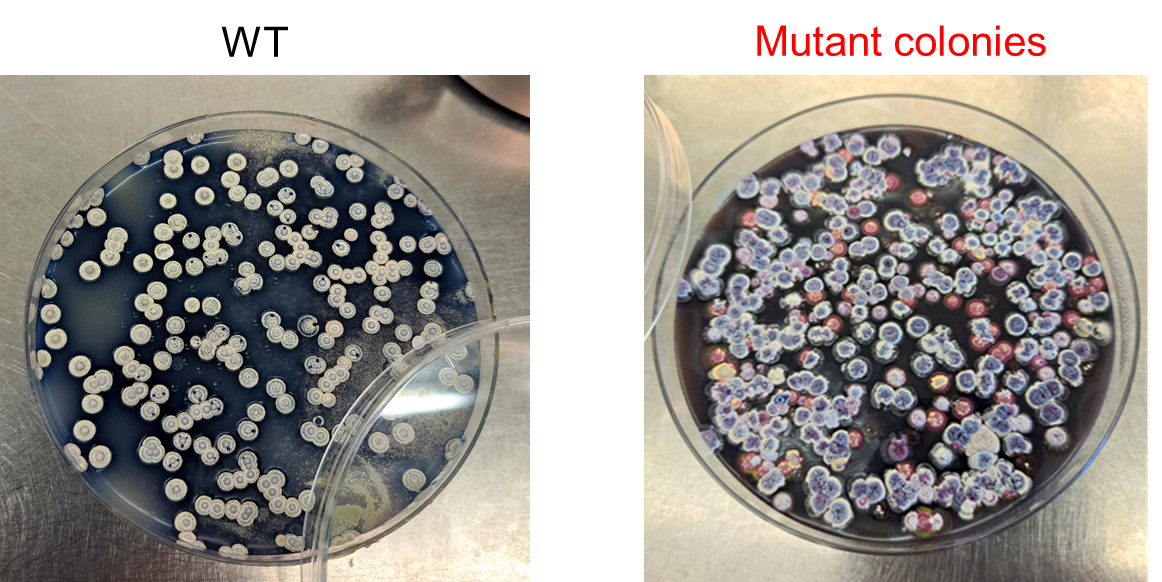

**Figure 1S**. 5 day old *Streptomyces coelicolor* colonies grown in SFM medium. Left image presents the WT morphology of *Streptomyces coelicolor,* white-gray and circular. Right image shows colonies with evident morphological changes of color and form obtained by replating a mutant colony.

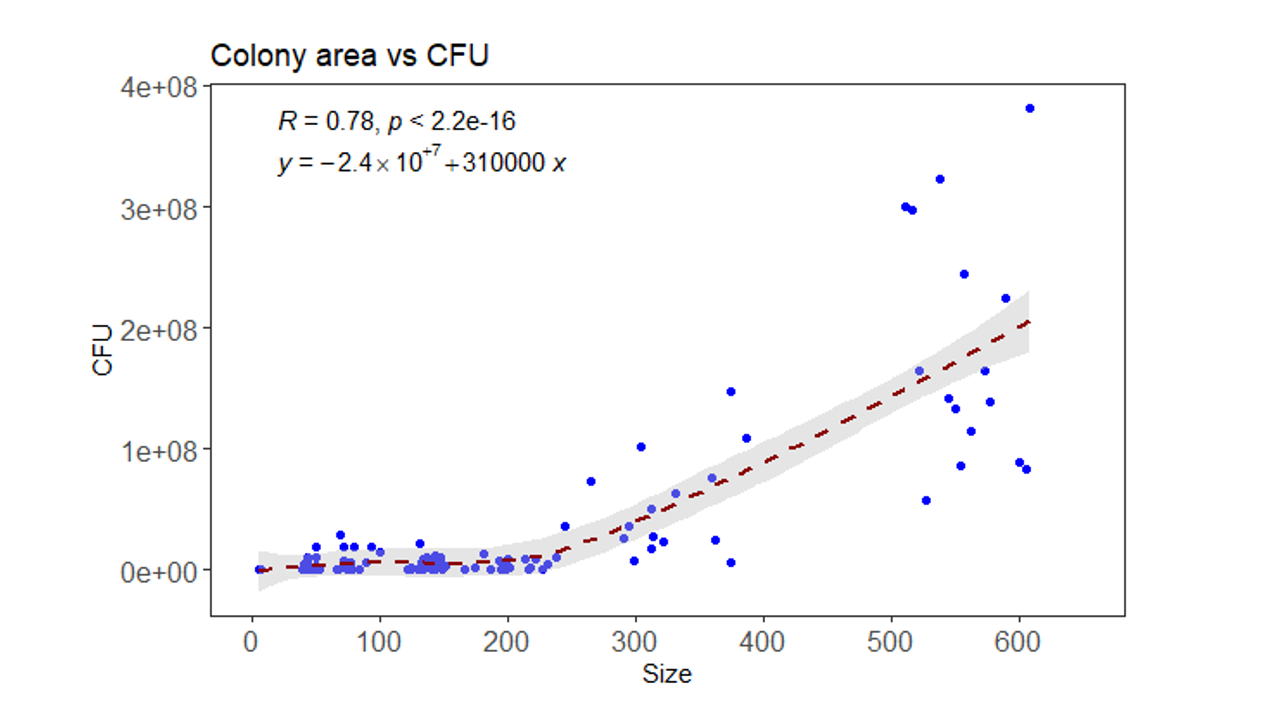

**Figue 2S.** Colony size with respect to CFU from colonies sampled through time.

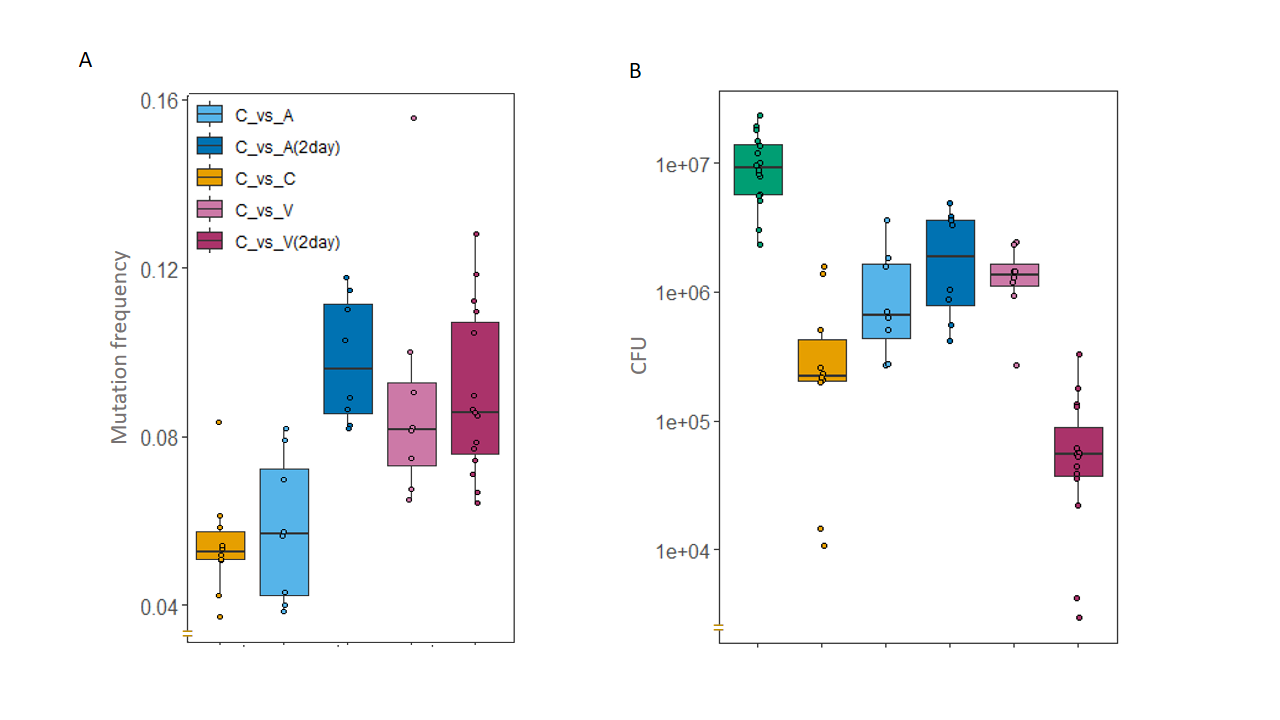

**Figure 3S** Competition between *Streptomyces coelicolor* and other species of *Streptomyces*. C = *S. coelicolor*, A = *Streptomyces ardus*, V = *S. venevuelae*. For strains with the label (2day) competitor *Streptomyces* specie was grown 2 days before.
